# Competition for H2A.Z between genes and repetitive elements establishes response to anti-viral immune activation

**DOI:** 10.1101/2022.03.31.486614

**Authors:** Fanju W Meng, Kristin E Murphy, Claire E Makowski, Patrick J Murphy

## Abstract

Activation of endogenous retroviruses leads to widespread transcriptional reprograming, affecting innate immune activation, metabolic control, and development. We find that the histone variant H2A.Z plays a central role in orchestrating these responses. Stimulating retroviral expression in zebrafish embryos causes H2A.Z to exit developmental gene promoters, which become silent, and to accumulate specifically at ‘primed’ repetitive elements, which are pre-marked by H3K27ac and H3K9me3. Remarkably, this rewiring is greatly influenced by total H2A.Z abundance, and developmental consequences of retrovirus activation are mitigated by H2A.Z over-expression. Our results uncover mechanisms whereby H2A.Z levels determine sensitivity to retroviral activation, and repetitive elements function as a nuclear sink to dramatically influence total transcriptional output.

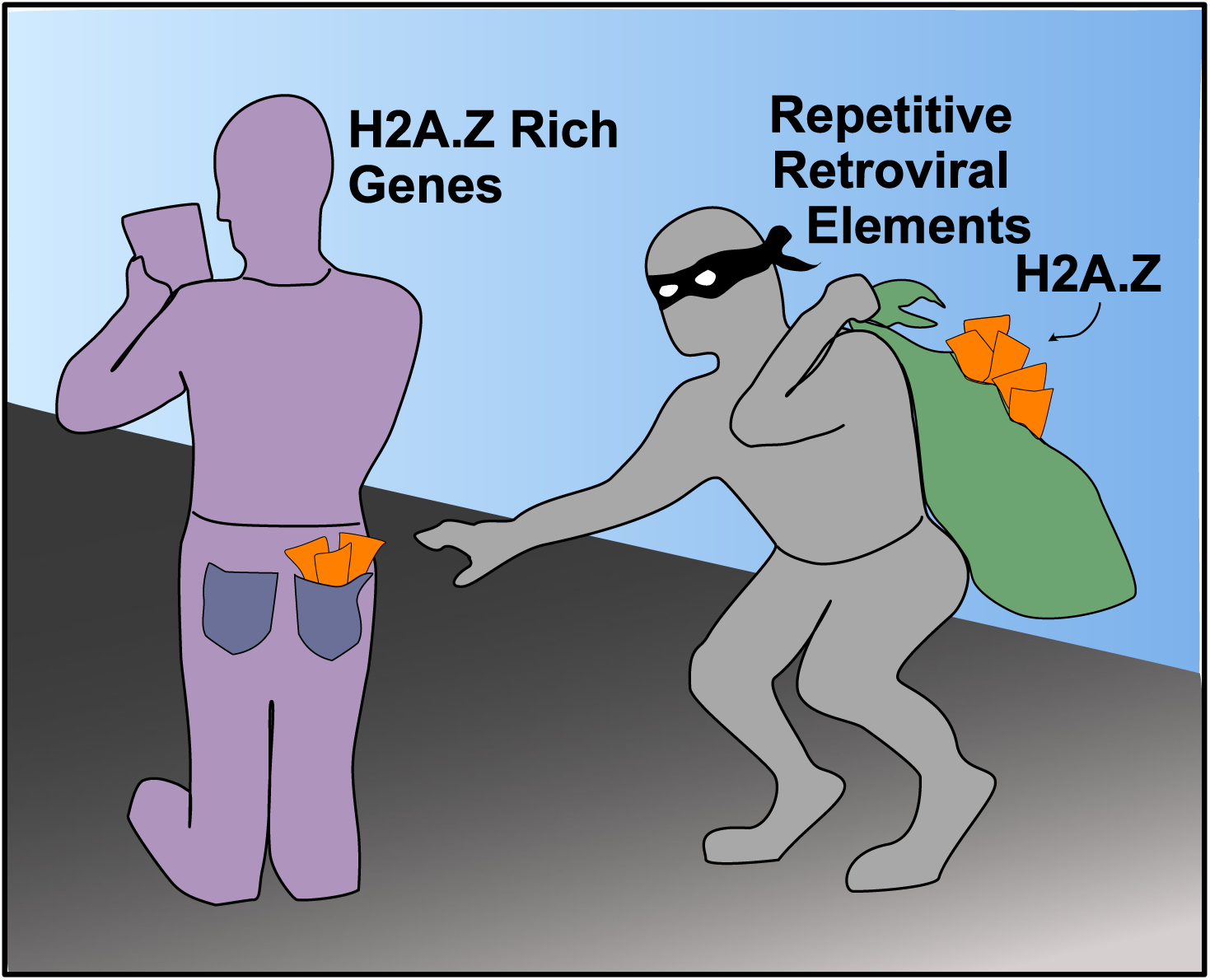

## Introduction

Large portions of eukaryotic genomes consist of ancestral repetitive elements (REs), including viral remnants from prior generations which can cause mutagenesis upon reactivation^1^. Several mechanisms have evolved for silencing REs^1^, including repressive epigenetic factors. Tri-methylation of histone H3 at lysine 9 (H3K9me3) and DNA methylation (DNAme) occur at the majority of REs and together these epigenetic marks promote transcriptional repression^2^. Recent studies have shown that removal of these marks causes some REs to become active^3^, but the majority of loci remain silent, and molecular features distinguishing expressed REs remain undetermined.

Evidence indicates that many evolutionarily younger REs can function as enhancers, and influence the expression of nearby genes upon reactivation^4^. Perhaps the best example for this occurs during viral infection response, when interferon induced transcription factors bind specifically at retroviral REs and regulate inflammation response genes^5^. In the absence of viral infection, activated REs can also trigger interferon signaling and inflammation response, mimicking viral infection through double stranded RNA synthesis^6,7^. Remarkably, discrete environmental stimuli and select cancer drugs have been shown to operate through this non-canonical axis, ultimately leading to reduced proliferation and apoptosis in exposed tissues or cancer cells^6,7^. Metabolism genes and cell-type specific gene pathways are often repressed as a consequence of RE activation^8^, but mechanisms linking primary effects, such as anti-viral response, to these common secondary effects, are lacking.

Recent studies have placed a renewed emphasis on histone variants as major players in transcriptional regulation of REs. The histone variant H3.3 recruits H3K9me3 at REs in mouse stem cells^9^, and studies from fruit flies indicate that the histone variant H2A.Z (h2av in flies) may act similarly^10^. H2A.Z resides predominantly at promoters surrounding the transcription start site (TSS) and has thus been studied primarily in the context of gene activation^11^. However, H2A.Z has also been detected at some mammalian REs, raising the possibility that it may function as a locus-specific silencing factor^11^. Despite these observations, a defined role for vertebrate H2A.Z in transcriptional control of REs has yet to been established.

## Main

### H2A.Z marks activate repetitive elements

Zebrafish are highly amenable to *in vivo* studies of retroviral function, innate immune activation, and epigenetic response to drug or toxicant exposure^12–15^. We therefore profiled shield stage zebrafish embryos using CUT&Tag^16^, in order to investigate H2A.Z function in control of RE activation (**Extended Data Fig.1a,b**). We found H2A.Z to be enrichment both at genic regulatory regions (**Fig.1a**), and at intergenic loci marked by H3K9me3 (**Fig.1b)**. These intergenic H3K9me3-marked loci contained all major classes of REs (**Extended Data Fig.1c**) and interestingly, they also possessed high levels of H3K27ac and DNAme (**Extended Data Fig.1d)**, epigenetic marks which rarely occur together elsewhere in the genome^17^.

**Fig.1:**
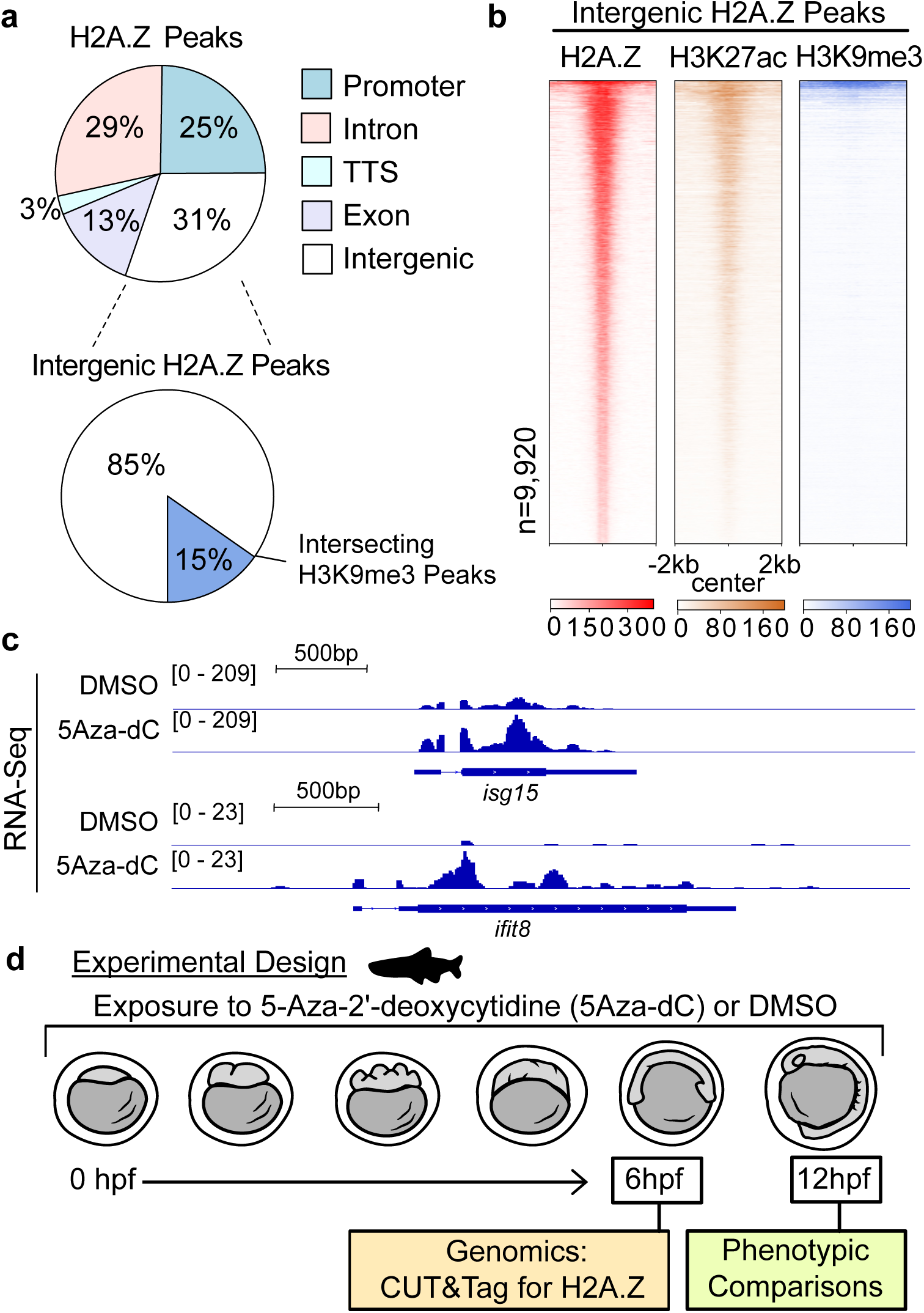
H2A.Z marks a subset of REs, coinciding with H3K27ac and H3K9me3. **a**, Genome annotation for H2A.Z peaks (top) and percent of intergenic H2A.Z intersection with H3K9me3 (bottom). **b**, Heatmaps of H2A.Z, H3K27ac and H3K9me3 over intergenic H2A.Z peaks. **c**, Genome browser snapshots showing transcription for *isg15* and *ifit8* in exposed embryos. **d**, Experimental design strategy.

The presence of H2A.Z at REs marked by both activating (H3K27ac) and silencing (H3K9me3 and DNAme) epigenetic marks led us to speculate that these REs may be temporarily maintained in a poised transcriptional state, such that they become active later during development, or in response to external stimuli/stressors. To investigate this possibility, zebrafish embryos were exposed to a known activator of RE expression, the DNA methyltransferase inhibitor 5-Aza-2’-deoxycytidine (5Aza-dC)^6,7^. As in prior studies, exposure from 1 to 6 hours post fertilization (hpf) caused anti-viral innate immune response genes to become active (**Fig.1c,d**), as well as RE activation (**Extended Data Fig.1e**). We next repeated H2A.Z CUT&Tag experiments under these 5Aza-dC exposure conditions, in order to better assess H2A.Z function in control of REs. DNAme covers the vast majority of the zebrafish genome in early embryos^18^, and based on the established antagonism between DNA methylation and H2A.Z binding^19–21^, we anticipated that removal of DNAme (resulting from 5Aza-dC treatment) would lead to increased H2A.Z enrichment throughout the genome. Instead however, genomic H2A.Z enrichment mostly decreased following 5Aza-dC exposure (**Fig.2a,b, Extended Data Fig.2a,b**). H2A.Z losses occurred primarily at the most highly enriched H2A.Z-marked promoters, which associated with transcriptional down-regulation (**Fig.2c**), and genes involved in metabolism were most affected (**Extended Data Fig.2c,d**). Notably, H2A.Z increases occurred primarily over REs, supporting the possibility that H2A.Z may be involved in their activation rather than silencing (**Fig.2d**).

**Fig.2:**
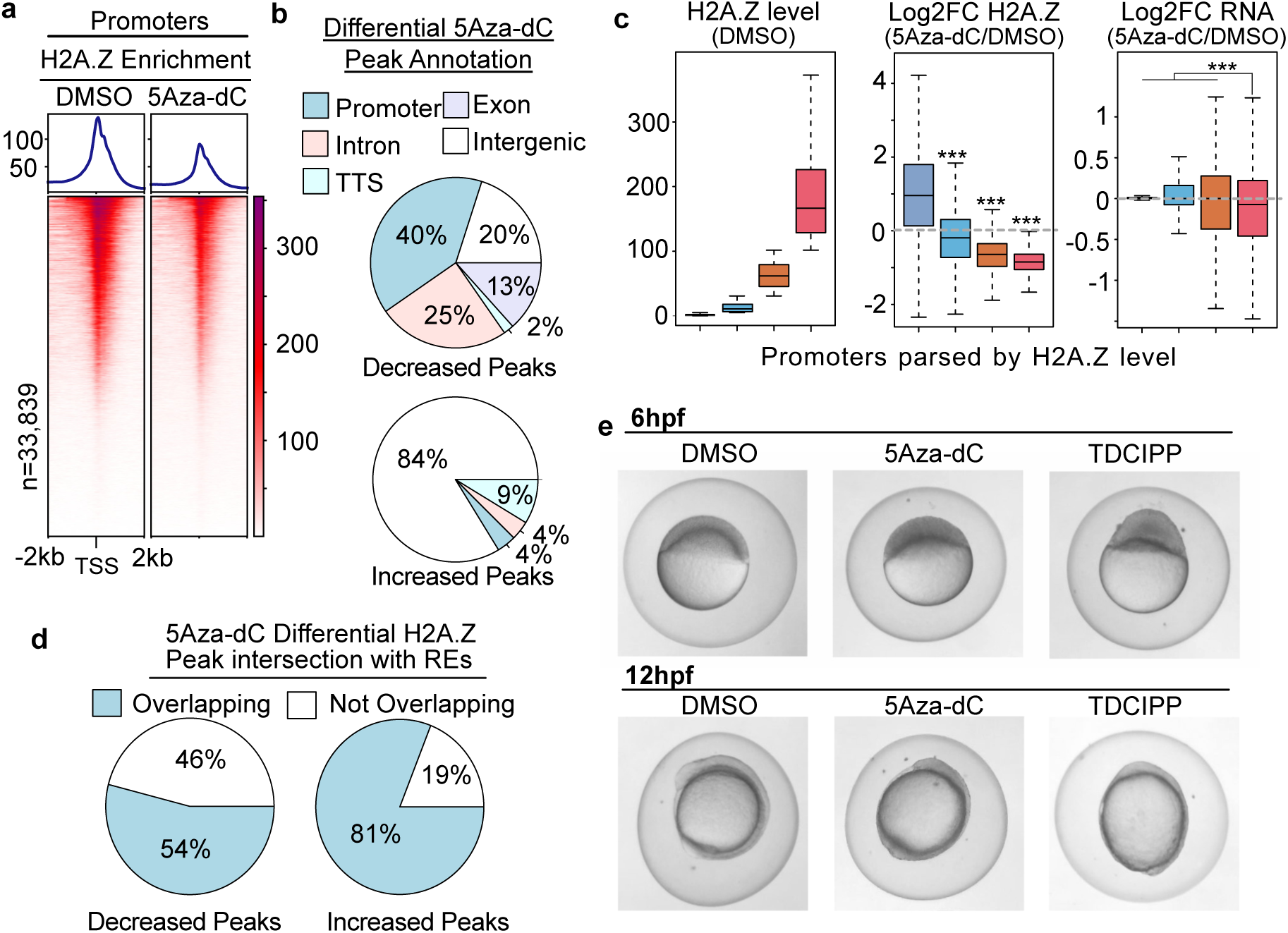
H2A.Z exits gene promoters and accumulates at REs following 5Aza-dC exposure. **a**, Heatmaps of H2A.Z enrichment over promoters (TSS+/-2kb, n=33,839) following 5Aza-dC exposure. **b**, Genome annotation for decreased (n=3,739) and increased (n=89) H2A.Z peaks. **c**, Boxplots for change in H2A.Z and gene expression, quartiles based on H2A.Z enrichment at 6hpf, *** = p-value < 0.001 from Student’s t-tests. **d**, Pie charts showing overlap between 5Aza-dC differential H2A.Z peaks and REs. **e**, Phenotypic comparisons between 5Aza-dC and TDCIPP treated embryos at 6 and 12hpf.

To investigate whether these epigenetic outcomes were unique to 5Aza-dC exposure, we next treated embryos with an additional environmental toxicant know to stimulate anti-viral innate immune activation^22,23^, the flame retardant Tris(1,3-dichloroisopropyl)phosphate (TDCIPP). TDCIPP exposure caused developmental defects similar to 5Aza-dC treatment (**Fig.2e**), along with like-wise H2A.Z reduction over promoters (**Extended Data Fig.3a-d**) and H2A.Z increases over REs (**Extended Data Fig.3d**). As was the case for 5Aza-dC exposure, H2A.Z losses corresponded with reduction in gene expression levels (**Extended Data Fig.3e**) and genes involved in metabolism were preferentially affected (**Extended Data Fig.3f**). In summary, induced reactivation of REs caused genome-wide re-localization of H2A.Z, with ectopic H2A.Z accumulating at REs and decreasing at promoters, which became silenced.

### Primed REs are predisposed to activation

Specificity in H2A.Z accumulation led us to speculate that H2A.Z might be involved in RE activation following 5Aza-dC treatment. In support of this hypothesis, H2A.Z enrichment was highest at REs which were initially active, and those which became active following 5Aza-dC treatment (**Fig.3a**). Activated REs were also enriched for H3K27ac and H3K9me3 (**Fig.3b**), epigenetic marks that typically function in opposition to one another. Aside from this unique combination of features, activated REs were epigenetically similar to other gene regulatory regions (including promoters and CpG islands) (**Extended Data Fig.4**). We therefore defined REs with the highest initial levels of H3K27ac and H3K9me3 as ‘primed’ (**Fig.3c)**, based on the fact that they were predisposed to both H2A.Z accumulation and activation following 5Aza-dC exposure. Primed REs included LINEs, LTRs, and tRNAs (**Fig.3d**), they tended to be evolutionarily younger (**Fig.3e**), and similar to enhancer elements, they were proximal to coding genes (**Fig.3f**) and enriched for transcription factor binding motifs (**Fig.3g**). These REs had high levels of H2A.Z initially (**Extended Data Fig.5a**), and levels increased further upon either 5Aza-dC or TDCIPP treatment (**Fig.3h, Extended Data Fig.5b**). Increases in H2A.Z enrichment were also accompanied by modest reduction in H3K9me3 levels (**Fig.3i, Extended Data Fig.5c)**, and sites with the greatest increase in H2A.Z tended to be those with the greatest reduction in H3K9me3 (**Extended Data Fig.5d,e)**. In sum, primed REs acquired H2A.Z, they became uniquely activated upon exposure to either 5Aza-dC or TDCIPP, and they possess several features of enhancers.

**Fig.3:**
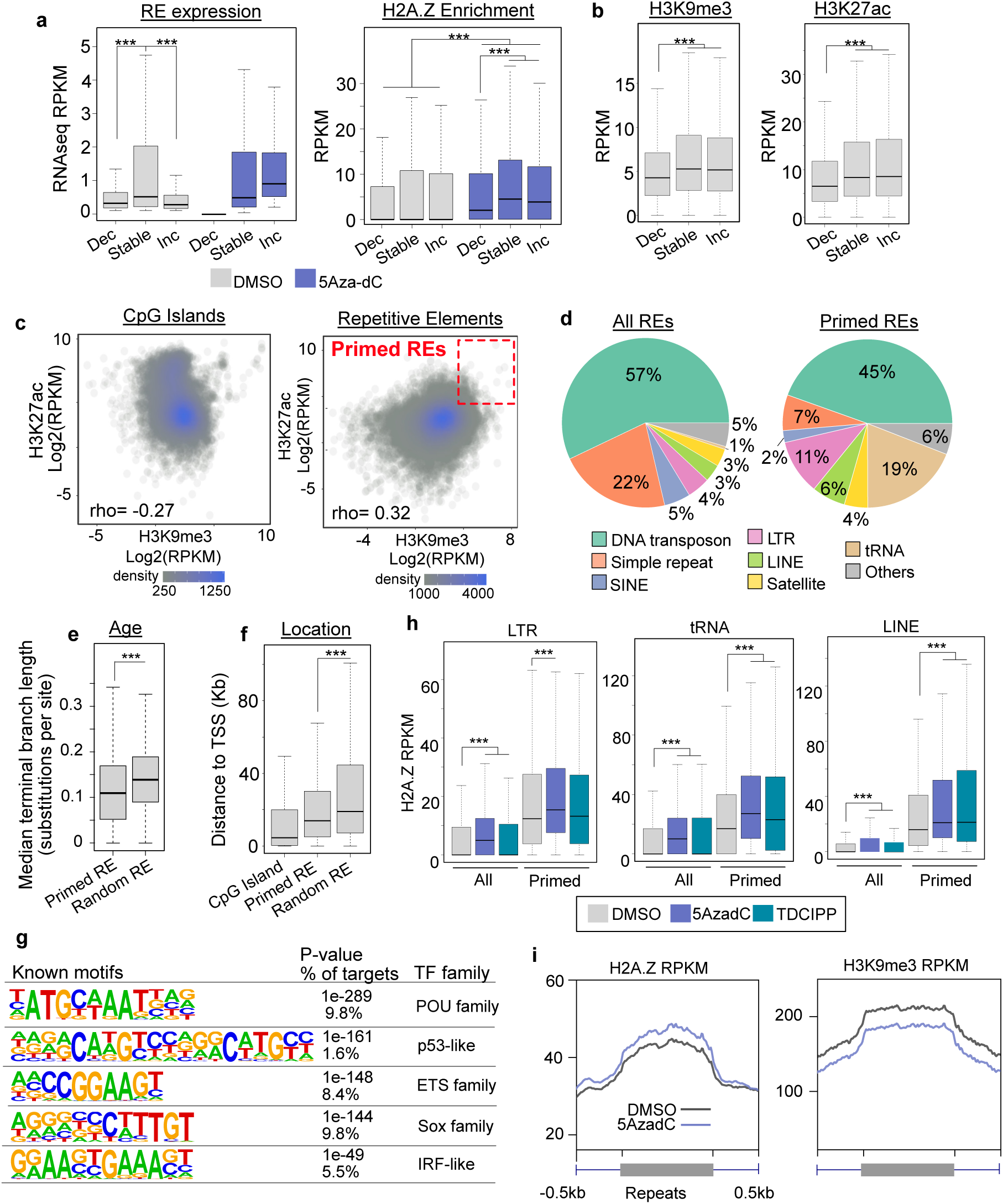
Primed REs exhibit unique chromatin features. **a**, Boxplots for transcription and H2A.Z enrichment at REs which decreased (Dec), increased (Inc) and remained transcriptionally stable following 5Aza-dC treatment. **b**, Boxplots of H3K9me3 (left) and H3K27ac (right) enrichment for RE groups as in a. **c**, Scatterplot of H3K9me3 and H3K27ac at CpG islands and REs. Statistics from Spearman’s correlation (primed REs, n=24,934). **d**, Portion of primed REs divided between major RE families. **e**, Boxplot for median evolutionary terminal branch length (substitutions per site) of primed and random REs. **f**, Boxplot for distance to nearest TSS comparing between genomic elements. **g**, Representative transcription factor motifs identified by Homer for primed repetitive elements. Non-primed repetitive elements were used as background control set. **h**, Boxplots for separate RE families quantifying H2A.Z enrichment following 5Aza-dC or TDCIPP exposure. **i**, Profile plots of H2A.Z and H3K9me3 changes at primed repeats following 5Aza-dC exposure. In a, b, e-h, statistical significance was determined using Student’s t-test, *** = p-value < 0.001.

### Competition for H2A.Z underlies response

H2A.Z reduction at promoters following 5Aza-dC exposure was accompanied by reduced gene expression levels (**Fig.2a-d**), and interestingly, many affected genes are dynamically expressed over normal developmental progression (**Extended Data Fig.6a**), suggesting that proper control of H2A.Z and/or DNAme may be critical for regulating developmental gene transcription. One compelling possibility is that total H2A.Z levels may have been insufficient in WT embryos to accommodate both developmental gene expression control and anti-viral response following 5Aza-dC exposure. Interestingly, average H2A.Z levels at primed REs are comparable to levels at enhancers (**Fig.4a**). These observations led us to hypothesize that focal H2A.Z losses at promoters might have occurred at the expense of H2A.Z accumulation at these activated REs. To investigate this possibility, we repeated 5Aza-dC treatment under conditions where H2A.Z was transgenically over-expressed from the *h2afx* promoter and marked by mCherry fluorescence^24^ (**Extended Data Fig.6b)**. As in wild type embryos, 5Aza-dC treatment caused innate immune response genes to become active (**Extended Data Fig.6c,d)**, and remarkably, embryos with increased levels of H2A.Z were partially resistant to 5Aza-dC treatment (**Fig.4b)**. We observed diminished morphological defects both at 12hpf and 24hpf. At 12hpf, growth of the head and extension of the tailbud were dramatically diminished by exposure to 5Aza-dC treatment, but over-expression of H2A.Z significantly rescued these defects (**Fig.4c,d**). At 24hpf, the majority (> 75%) of WT zebrafish embryos exposed to 5Aza-dC displayed severe morphological defects, but in the presence of H2A.Z over-expression, severe defects were far less common (< 25%), and many embryos appeared phenotypically normal or were only moderately impacted (**Fig.4e,f**).

**Fig.4:**
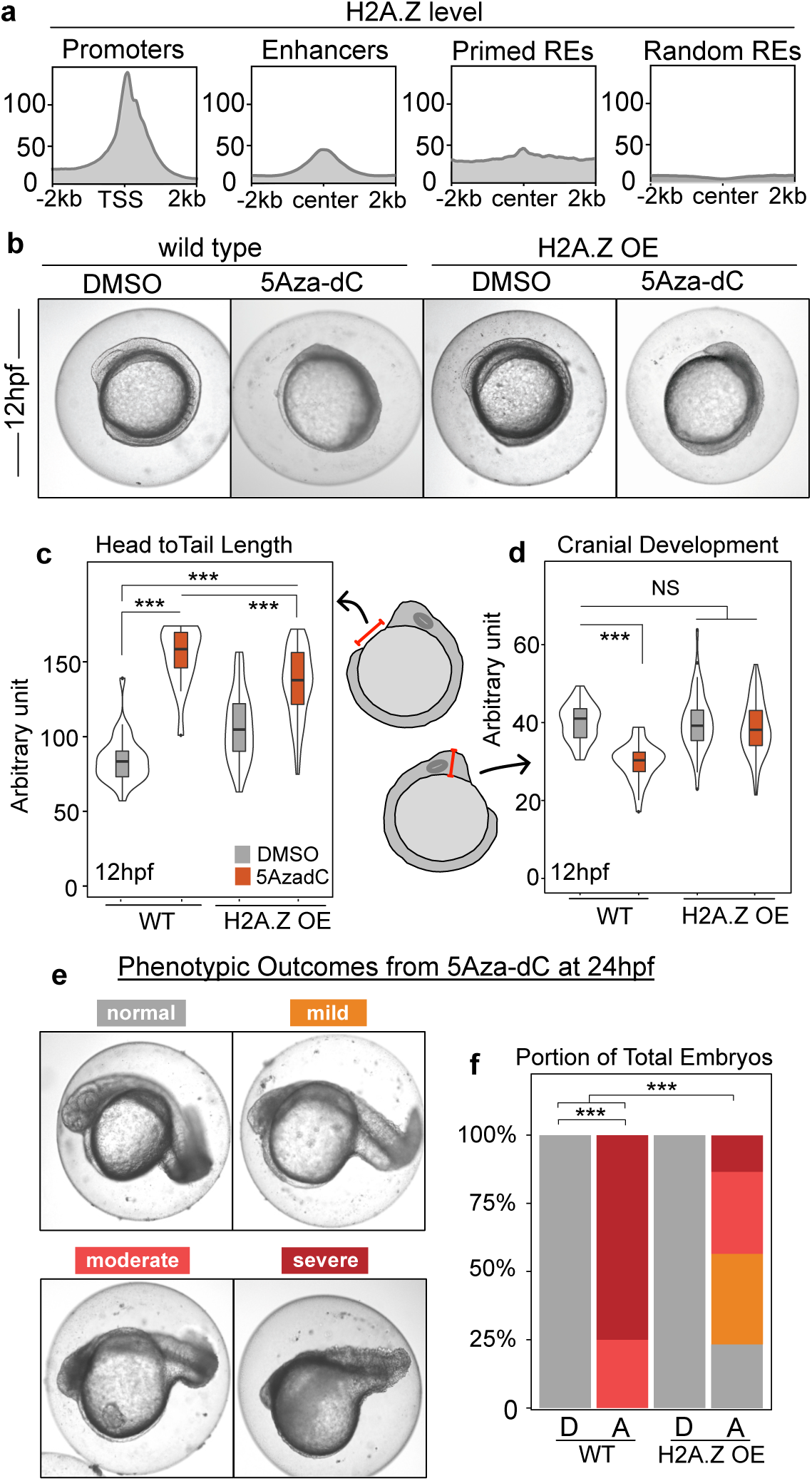
H2A.Z overexpression mitigates 5Aza-dC induced developmental defects. **a**, Profile plots of H2A.Z signals at promoters, putative enhancers, primed and random REs. **b**, Representative images of 12hpf wild type and H2A.Z^mCherry^ (OE, overexpression) embryos following 5Aza-dC treatment. **c**-**d**, Violin boxplot for the length between head and tail (c) and maximum length of head region (d) in wild type and H2A.Z^mCherry^ embryos treated with 5Aza-dC at 12hpf. **e**-**f**, Representative images for phenotypic classes in embryos at 24hpf and quantifications of these classes in wild type or H2A.Z^mCherry^ embryos following treatment [A = 5Aza-dC, D = DMSO] Fisher’s exact test, *** = p-value < 0.001.

To investigate the degree to which 5Aza-dC induced gene expression changes were mitigated by H2A.Z over-expression, we next performed whole embryo transcriptome analysis of exposed WT and H2A.Z-mCherry embryos at 12hpf (**Fig.5a)**. Similar to our findings at 6hpf (**Extended Data Fig.6a)**, genes dysregulated by 5Aza-dC exposure normally become active at slightly later stages of development (**Extended Data Fig.6e,f**), and in agreement with the observed phenotypic outcomes, over-expression of H2A.Z partially reversed impacts on gene expression (**Fig.5a-c**). Transcriptional changes for both up- and down-regulated genes were partially reversed in the presence of H2A.Z-mCherry (**Fig.5b,c**). Gene ontology analysis revealed that down-regulated genes tended to function in developmental progression, whereas up-regulated genes tended to function in stress response, apoptosis, and cell death (**Fig.5d)**. Taken together, these results demonstrate that total H2A.Z levels greatly influence the developmental defects and transcriptome-wide dysregulation that occurs following exposure to 5Aza-dC.

**Fig.5:**
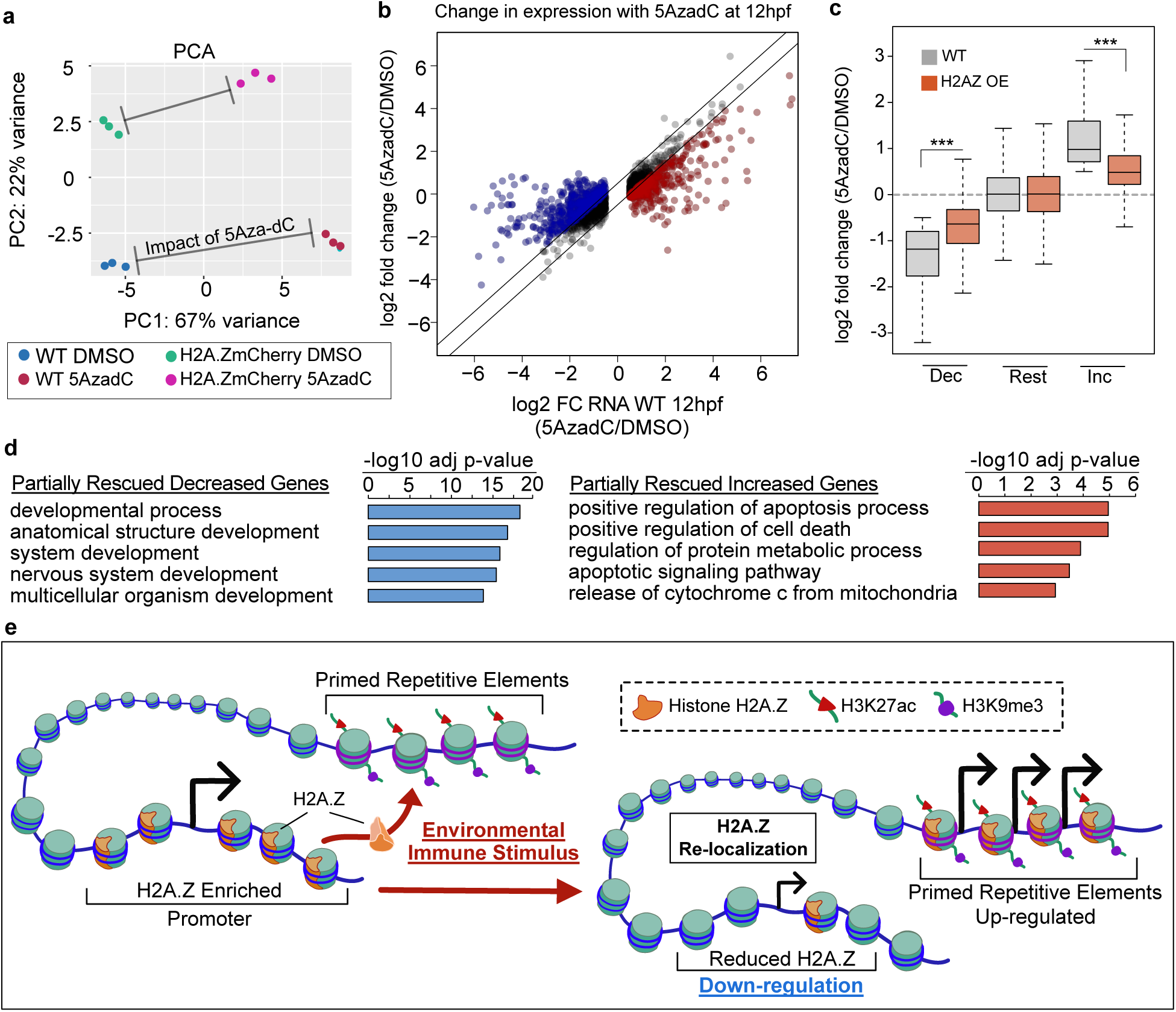
H2A.Z overexpression partially rescues 5Aza-dC induced transcriptional changes. **a**, Principal component analysis (PCA) of RNA-seq replicates of DMSO and 5Aza-dC treated wild type and H2A.Z^mCherry^ embryos at 12hpf. **b**, Scatterplot of log2 fold change of gene expression (5Aza-dC/DMSO) for wild type and H2A.Z^mCherry^ embryos at 12hpf. Partially rescued genes in H2A.Z^mCherry^ embryos were colored. **c**, Boxplot for log2FC (5Aza-dC/DMSO) in gene expression for wild type (gray) and H2A.Z^mCherry^ embryos (pink) at 12hpf following 5Aza-dC treatment. Genes separated by impact after 5Aza-dC in wild type [Dec = decreased, Inc = increased, Rest = remaining]. Statistical significance was determined using Student’s t-test, *** = p-value < 0.001. **d**, Enriched gene ontology terms associated with those partially rescued decreased and increased genes as defined in b. **e**, Model depicts sequestration of H2A.Z at primed REs in response to RE activation and consequential impacts on genes that are normally active.

## Discussion

Results of this study demonstrate that differences in H2A.Z abundance significantly influence how embryos are impacted by RE activation, with low H2A.Z levels conferring sensitivity and high levels conferring robustness/resistance. In this sense, individual organisms may be more or less sensitive to retro-viral activation dependent on initial H2A.Z levels. Interestingly, prior studies have found H2A.Z to be over-expressed in many forms of cancer^25^, perhaps providing tumors with a means to evade cell death following innate immune activation. In this regard, targeting H2A.Z or regulators of H2A.Z installation may uncover effective strategies to combat cancer, especially in the context of retro-viral activation, or in tumor that are resistant to viral-mimicry-based therapies^26^.

Examination of genes dysregulated by 5Aza-dC suggests that the mechanisms we describe here may contribute to the gene expression changes during normal developmental progression. For both 6hpf and 12hpf embryos, genes dysregulated by 5Aza-dC exposure corresponded with those that are normally activated at distinct developmental timepoints. These results are reminiscent of studies in mammals, where activation of enhancer-like endogenous retroviruses promotes embryonic genome activation and pluripotency^27–29^, or of neural crest development in zebrafish, where the neural crest marker gene *crestin* is derived from an RE^30^. Prior studies in zebrafish indicate that similar REs may even function in the establishment of higher-order chromatin structure during embryonic development^31^. Evidently, further research is necessary to investigate the role that REs may have during vertebrate developmental transitions and/or the emergence of particular lineages.

We found the combined presence of H3K27ac and H3K9me3 at active REs as well as those which became active upon exposure to 5Aza-dC or TDCIPP. Based on the opposing nature of these histone marks, we hypothesize that they may function in a manner analogous to bivalent chromatin domains^32^, such that particular REs are ‘primed’ or ‘predisposed’ to future activation. Interestingly, primed REs tended to be evolutionarily younger, as compared with typical REs, indicating that they may have arisen from recent transposition at already active/accessible loci^33^, or perhaps ‘primed’ REs have maintained the ability to become transcriptionally active despite counteractive silencing mechanisms. Regardless of origin, longitudinal studies are necessary to test whether epigenetic ‘priming’ at REs confers increased sensitivity during toxicant exposure, pharmacological therapy, or stress response.

The mechanism we present here, whereby H2A.Z is sequestered to particular REs upon their activation (**Fig.5e**), may represent a general principle in eukaryotic biology, and although we focus specifically on H2A.Z, it is certainly possible that other nuclear factors are sequestered in similar manner. In this regard, our study presents a conceptual advance in the broader understanding of transcription control, whereby repetitive DNA sequences and cell-type-specific genes networks engage in an intrinsic competition for transcription machinery, in effect governing overall cellular output while simultaneously enabling response to external stimuli.

## Methods

### Animals and chemicals

Adult wild type Tübingen and H2A.Z^mCherry^ (*Tg(h2ax:h2az2a-mCherry*, ZL1953)) zebrafish were maintained on a 14h:10h light:dark cycle. Newly fertilized eggs were used for all experiments performed in this study. Zebrafish husbandry and care were conducted in full accordance with animal care and use guidelines with ethical approval by the University Committee on Animal Resources at the University of Rochester Medical Center. 5-aza-2’-deoxycytidine (5Aza-dC) was purchased from Cayman Chemical (Item No. 11166, Purity > 98%), TDCIPP was purchased from MedChemExpress (Cat. No HY-108712, Purity > 98%), and DMSO were purchased from VWR Life Science.

### Embryonic treatment of 5Aza-dC and TDCIPP

Newly fertilized zebrafish eggs were collected after spawning, and approximately 50 embryos per well were maintained in a 6 well plate within a light and temperature-controlled incubator. As in prior studies, embryos were treated in triplicates with 5 mL of 3.12 uM TDCIPP (treatment starts from 2-cell stage at 0.75 hour post fertilization), or 100 uM of 5Aza-dC (treatment starts from 1-cell stage), with 0.1% DMSO used for control group, and incubated until 6hpf (hour post fertilization) for H2A.Z CUT&Tag and 12hpf for RNA-seq, or 6hpf, 12hpf and 24hpf for morphological characterizations.

### Phenotypic characterization and quantification

To quantify phenotypic outcomes of 5Aza-dC and TDCIPP exposure, images of treated zebrafish embryos were captured at 6 hpf, 12 hpf or 24 hpf using levenhuklite software package with bench-top stereo microscope.

To compare phenotypes of 5Aza-dC treated wild type and H2A.Z^mCherry^ embryos at 12hpf, lateral images were taken for each embryo at the same setting, and a straight line was drawn and measured in ImageJ (version 2.0.1/1.53c) using “Straight” tool between head and tail, and between edge of head and yolk sac for maximum head width. Violin and boxplots were used to present the measures for wild type and H2A.Z^mCherry^ embryos treated with DMSO and 5Aza-dC.

To compare phenotypes of 5Aza-dC treated wild type and H2A.Z^mCherry^ embryos at 24hpf, embryos were scored into four groups based on severity of morphological defects. Normal: embryos with well developed head and normal length of tail; mild: embryos with well developed head and slightly shortened length of tail; moderate, embryos with poorly developed head and obviously shortened tail; severe, embryos with barely developed head and dramatically shortened tail.

### Genomic H2A.Z and H3K9me3 profiling by Cleave Under Targets and Tagmentation (CUT&Tag)

Treated embryos were collected at 6 hpf, and manually dechorionated. Then, embryos were transferred into 1.5ml Eppendorf tubes, and dissociated by pipetting. Cells were pelleted after centrifugation at 600g for 3 minutes, and nuclei were extracted on ice for 10 minutes, and collected by centrifugation at 600g for 3 minutes. Around 100,000 nuclei were used for each replicate, and CUT&Tag libraries were generated according to published CUT&Tag protocol^1^. Polyclonal H2A.Z antibody (Active motif, Cat. No 39113, 1:50 dilution) and H3K9me3 antibody (Active motif, Cat. No 39062, 1:50 dilution) were used for CUT&Tag.

### CUT&Tag libraries sequencing and data analysis

CUT&Tag libraries were pooled and sequenced using Illumina paired-end sequencing HiSeq 4000 platform. Raw sequencing reads were checked for quality using FastQC (version 0.11.9), and adapter sequence were removed using Cutadapt (version 2.7). Sequencing reads were aligned to zebrafish genome assembly (GRCz11) using Bowtie2 (version 2.2.5) with default setting. Sequence duplicates were removed using Picard MarkDuplicates (version 2.5.0), and genome browser bigwig files for H2A.Z and H3K9me3 were generated using merged replicate bam files and converted to bigwig files using bamCoverage (version 3.5.1, --normalizeUsing RPKM --binSize 10). Integrative Genomics Viewer (version 2.8.13) was used to visualize all sequencing data and to prepare genome browser snapshots.

To identify H2A.Z occupied genomic regions, peak calling of H2A.Z CUT&Tag was performed using macs2 callpeak (version 2.2.6) with the following parameters: -f BAMPE -g 1.5e9 –nomodel --broad. Genomic annotation of H2A.Z peaks was performed using homer annotatePeaks.pl to assess annotations, distance to nearest promoters and CpG content. Bedtools intersect (version 2.30.0) was used to examine overlapping between intergenic H2A.Z peaks and H3K9me3 ChIP-seq peaks.

To identify differential H2A.Z peaks between DMSO and 5Aza-dC or TDCIPP treated embryos, H2A.Z broadPeak and associated bam files for each replicate were used as input files for DiffBind (version 3.2.5) with default setting. Significantly increased H2A.Z peaks were defined as adjusted p-value < 0.05 and log2 fold change > 0.5, while significantly decreased H2A.Z peaks were defined as adjusted p-value < 0.05 and log2 fold change < -0.5.

### Differential gene expression analysis of wild type and H2A.Z^mCherry^ embryos

DMSO and 5Aza-dC treated wild type and mCherry-positive embryos at 12hpf were used to extract RNA. Chorions were removed with forceps, and embryos were collected in Trizol and grinded with mortar and pestle. Total RNA was extracted with phenol:chloroform:isoamyl alcohol, then aqueous layer were purified with Zymo Direct-zol RNA Miniprep kit.

Total RNA-seq libraries were prepared and sequenced with paired-end 150bp reads by LC Sciences. Sequencing reads were mapped with STAR with default setting (v2.7.2a), and mapped reads were used in featureCounts (v2.0.3 in subread) using GRCz11.102.gtf. Count table was used to generate principal component analysis (PCA) plot by DESeq2 (v1.22.2 in R/3.5.1). WT 12hpf 5Aza-dC significantly increased genes were defined as adjusted p-value < 0.05 and log2 fold change > 0.5 (n=841), significantly decreased genes were defined as adjusted p-value < 0.05 and log2 fold change < -0.5 (n=1,002).

To examine changes in gene expression after 5Aza-dC or TDCIPP exposure at 6hpf, featureCounts (v2.0.3 in subread) was used to calculate reads over exons. Read counts were normalized to sum of read counts of each sample, and log2 fold change was calculated in R. Differentially expressed genes were identified based on log2 fold change: log2 fold change > 0.5 for increased genes, and log2 fold change < -0.5 for decreased genes.

To identify partially rescued genes in 5Aza-dC treated H2A.Z^mCherry^ embryos at 12hpf, differentially expressed genes in 5Aza-dC treated wild type embryos were identified using DESeq2 (log2FC at 0.5, and adjusted p-value < 0.05). Among those genes, partially rescued increased genes were those which have less than 0.5 difference in log2FC in H2A.Z^mCherry^ embryos compared to wild type embryos (log2FC_H2AZ^mCherry^ - log2FC_wildtype < 0.5). Similarly, partially rescued decreased genes were those more than 0.5 difference in log2FC in H2A.Z^mCherry^ embryos (log2FC_H2AZ^mCherry^ - log2FC_wildtype > 0.5).

### Developmental stage gene expression and gene set enrichment analysis (GSEA)

Read counts normalized (TPM) gene expression table from 1-cell stage to day 5 was publicly available^2^. To examine temporal dynamics of gene expression, particular gene sets were merged with count table in R to calculate average expression profile for plotting in R, and to generate gene expression heatmap in R using pheatmap.

GSEA was performed to determine whether any gene set of particular developmental stage was enriched in 6hpf and 12hpf 5Aza-dC treated wild type RNA-seq data, stage-specific maximum expressed genes were identified in R by assigning each gene to a developmental stage where maximum gene expression was scored using TPM count table from White *et al*. eLife, 2017. These stage-specific gene sets were used with GSEA, along with featureCounts table of 6hpf and 12hpf RNA-seq data to run GSEA (v4.1.0).

### Gene ontology analysis of differential peaks and genes\

Differential H2A.Z peaks and gene groups were used as input files, and enriched gene ontology terms associated with these regions and genes were identified by g:Profiler^3^. Significance threshold was set at 0.05 using g:SCS threshold for multiple testing.

### Identification of primed repetitive elements

To identify repetitive elements with high H3K9me3 and H3K27ac, bed file of all annotated zebrafish repeats (genome assembly GRCz11) was downloaded from UCSC table browser, and used as input bed file to calculate enrichment of H3K9me3 and H3K27ac using multibigwigsummary (version 3.5.1). Output matrix files were analyzed in R, and primed repeats were identified as enrichment score of top 5% for both H3K9me3 and H3K27ac. For comparisons, random genomic regions were identified using bedtools random -l 1000 -n 10000. Random non-primed repeats were randomly selected using R. To estimate evolutionary age of repetitive elements, each repeat was matched to its evolutionary age^4^. Median terminal branch length (substitutions per site) on phylogenetic tree were used for plotting. To identify enriched transcription factor binding motifs in primed repetitive elements, bed files for primed repetitive elements was used as input to HOMER findMotifsGenome.pl. “-size given” was used, and random non-primed repeats were used as background regions for motif discovery.

### Percentage quantification of repetitive elements

To quantify percentage of repetitive elements, repeats within the same superfamilies were counted, including DNA transposon, LTR, LINE, SINE, tRNA, Satellite DNAs and Simple repeats. Pie charts were generated using R (version 4.0.3).

### Transcription of repetitive elements loci

To quantify changes in transcription at repetitive element loci after 5Aza-dC treatment, RPKM normalized scores were calculated at each repeat using bigwig files of DMSO and 5Aza-dC RNA-seq data. Repeats with DMSO RPKM score less than 0.1 were excluded from downstream analysis. Repeats with potentially increased transcription were identified based on log2 fold change (5Aza-dC/DMSO) > 1 (Inc), and repeats with potentially decreased transcription were identified based on log2 fold change (5Aza-dC/DMSO) < -1 (Dec), all remaining repeats were grouped as ‘Stable’.

### Data representation

Heatmaps and profile plots were generated using plotHeatmap and plotProfile from deepTools (version 3.5.1) or pheatmap in R, and fonts and labels were adjusted in Affinity Designer (version 1.9.1). Density scatterplots were generated using ggplot2 geom_density. All boxplots were plotted using R, horizontal line shows the median, and the box encompasses the interquartile range.

## Data Availability

All sequencing datasets generated in this study were deposited to GEO data repository (GSE162207). Previously published genomics data that were used here are available under the following accession codes: GSE113086 (H3K9me3 ChIP-seq)^5^; GSE114954 (H3K27ac ChIP-seq)^6^; GSE130944 (ATAC-Seq)^7^; SRP020008 (DNA methylation and RNAseq of 5Aza-dC)^8^; BioProject ID PRJNA475635 (RNA-seq of TDCIPP)^9^. Normalized gene expression read counts table (TPM) across developmental stages from 1-cell to 5 dpf was publicly available^2^.

## Extended data figures

**Extended Data Fig.1.**
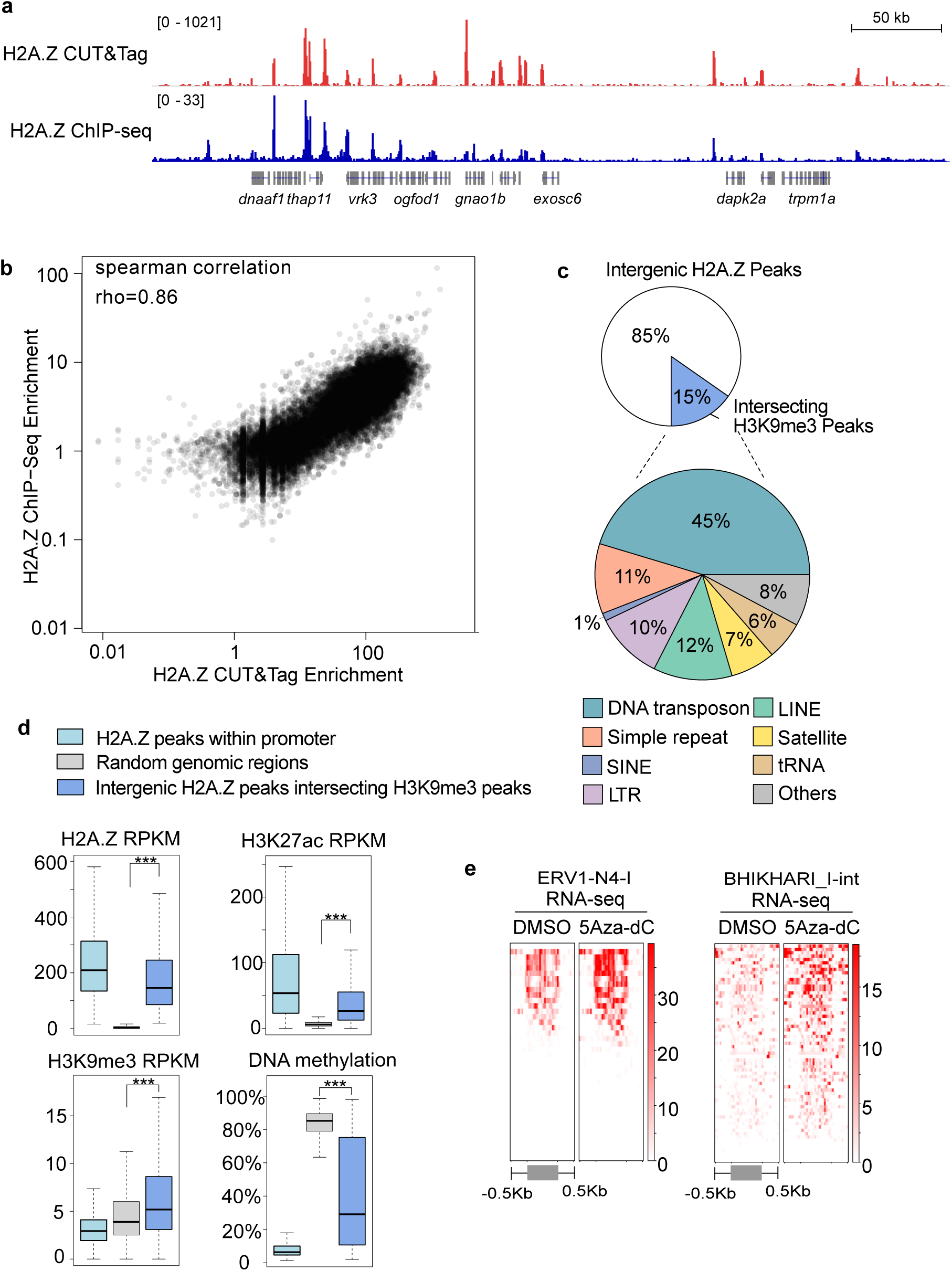
Comparison of H2A.Z ChIP-seq with H2A.Z CUT&Tag. **a**, Representative genome browser views demonstrate that H2A.Z CUT&Tag recapitulates H2A.Z genomic profiles established by H2A.Z ChIP-seq. **b**, Scatterplot showing high correlation (rho=0.86, spearman correlation) between H2A.Z CUT&Tag and H2A.Z ChIP-seq signals over gene promoter regions (TSS+/-1kb). **c**, Pie chart representing repetitive elements superfamilies that overlap intergenic H2A.Z peaks interesting H3K9me3 peaks. **d**, Boxplots of RPKM values for H2A.Z, H3K27ac, H3K9me3, and DNA methylation for H2A.Z peaks within promoter (cyan, n=7,963), random genomic regions (gray, n=7,582) and intergenic H2A.Z peaks intersecting H3K9me3 peaks (blue, n=8,018). Statistical significance was determined using Student’s t-test, ***, p-value < 0.001. **e**, Heatmaps of normalized RNA-seq reads over ERV1-N4-I and BHIKHARI_I-int repeats. Repeat lengths scaled into 1kb together with 0.5kb both upstream and downstream were used for heatmaps.

**Extended Data Fig.2.**
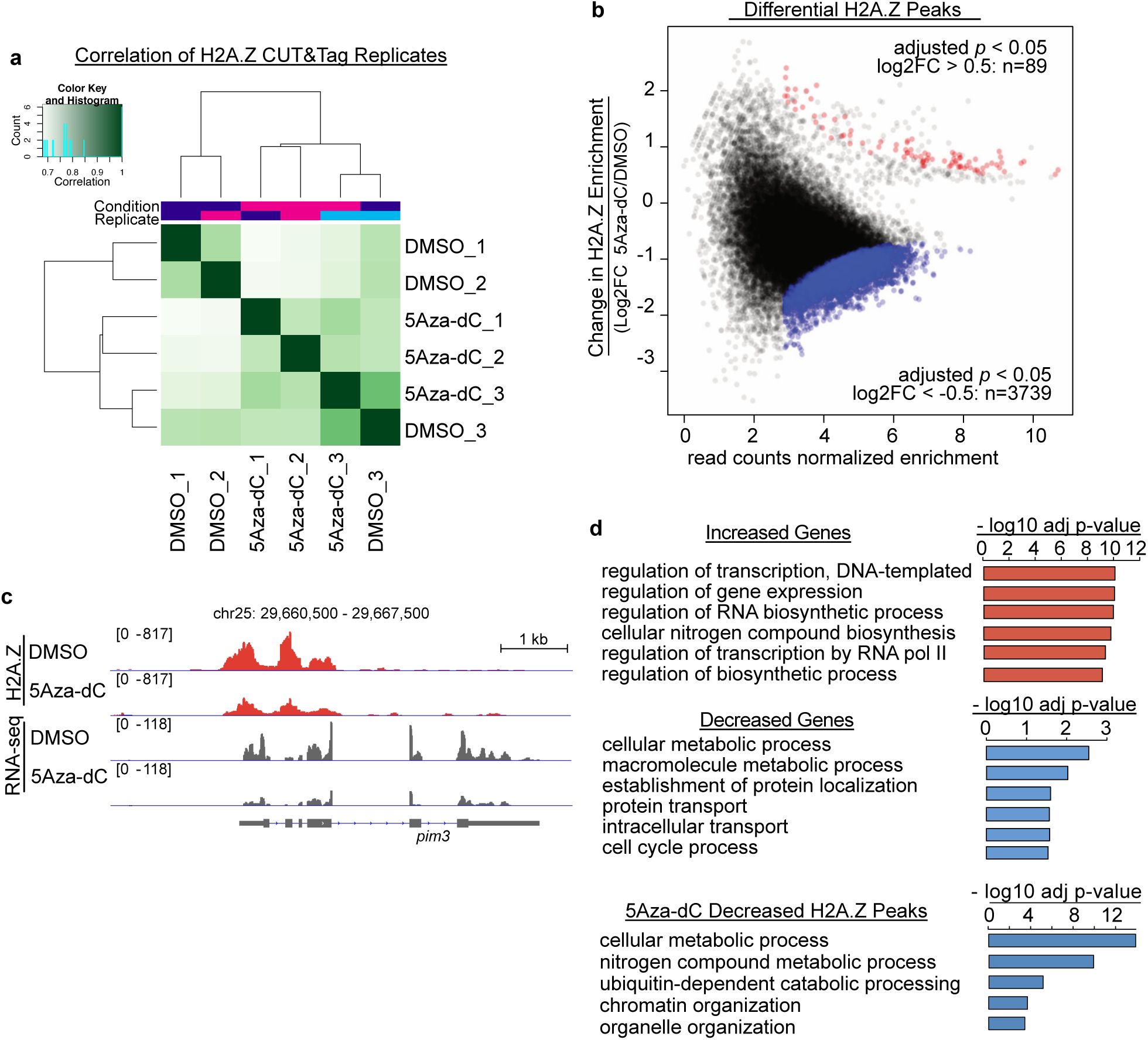
Changes of H2A.Z patterning in 5Aza-dC treated embryos. **a**, Correlation heatmap generated by DiffBind using read count of H2A.Z CUT&Tag samples from DMSO and 5Aza-dC treated 6hpf embryos. **b**, MA plot depicts H2A.Z peaks with differential H2A.Z levels after 5Aza-dC exposure in embryos at 6hpf. 89 regions were identified with significantly increased H2A.Z (shown in red, adjusted p-value < 0.05 & log2FC > 0.5) while 3,739 regions with significantly decreased H2A.Z (shown in blue, adjusted p-value < 0.05 & log2FC < - 0.5). **c**, Genome browser views of changes of H2A.Z and gene expression at the *pim3* locus. **d**, Enriched gene ontology terms for differential genes and H2A.Z peaks in 5Aza-dC treated 6hpf embryos. No enriched terms found for increased H2A.Z peaks.

**Extended Data Fig.3.**
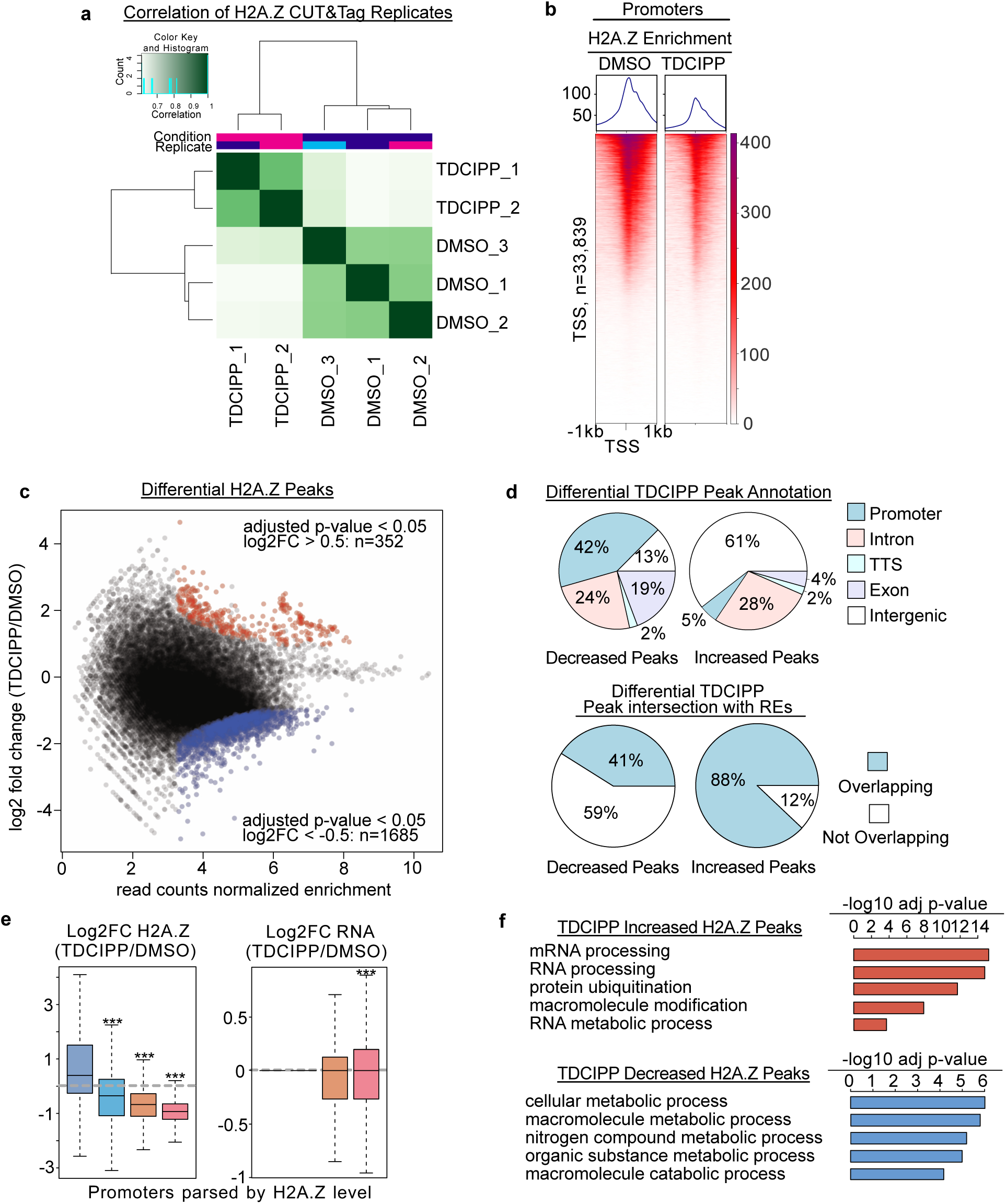
Changes of H2A.Z patterning after TDCIPP exposure. **a**, Correlation heatmap generated by DiffBind using read count of H2A.Z CUT&Tag samples from DMSO and TDCIPP treated embryos. **b**, Heat maps showing reduced H2A.Z at promoter regions in TDCIPP treated embryos. All heatmaps were ranked in the same order. **c**, MA plot depicts H2A.Z peaks with differential H2A.Z levels after TDCIPP exposure in 6hpf embryos. 352 regions were identified with significantly increased H2A.Z (shown in red, adjusted p-value < 0.05 & log2FC > 0.5) while 1,685 regions with significantly decreased H2A.Z (shown in blue, adjusted p-value < 0.05 & log2FC < -0.5). **d**, Pie charts represent genomic annotations of differential H2A.Z peaks (top) in TDCIPP treated embryos, and the degree of overlapping between TDCIPP differential H2A.Z peaks and repetitive elements (bottom). **e**, Boxplots of log2 fold change of H2A.Z and gene expression for promoters parsed into 4 groups based on H2A.Z levels in DMSO treated 6hpf embryos. Statistical significance was determined using two samples and one sample Student’s t-test (u=0), ***, p-value < 0.001. **f**, Enriched gene ontology terms for differential H2A.Z peaks in TDCIPP treated embryos.

**Extended Data Fig.4.**
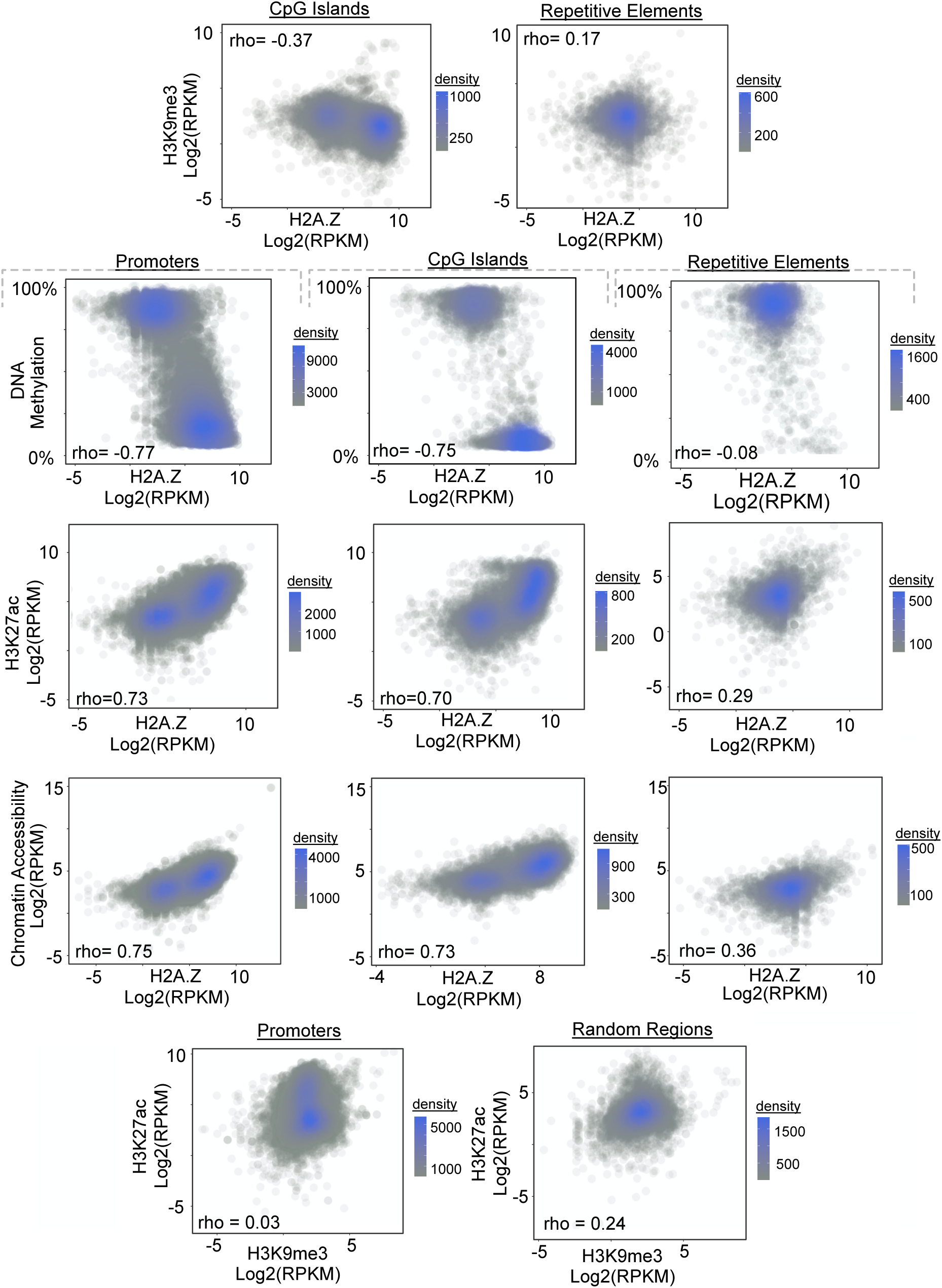
Primed repetitive elements exhibit unique chromatin features. Density scatterplots of H3K9me3, DNA methylation, H3K27ac, chromatin accessibility and H2A.Z for CpG islands, promoters, repetitive elements and random regions. Spearman’s correlation coefficients were shown.

**Extended Data Fig.5.**
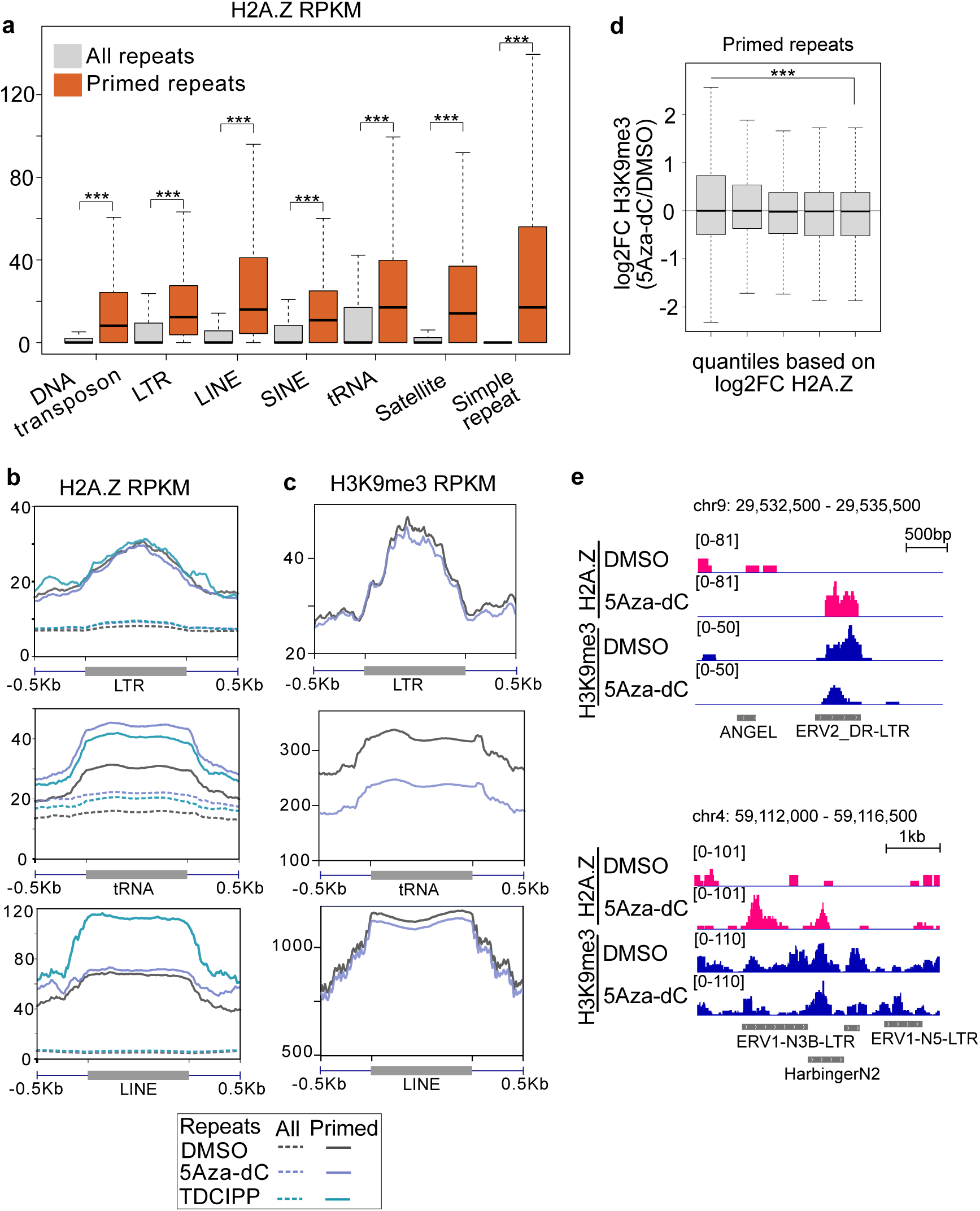
H2A.Z increases at primed repetitive elements. **a**, Boxplot of H2A.Z score of all repeats (gray) and primed repeats (orange) for major superfamilies. Statistical significance was determined using Student’s t-test, ***, p-value < 0.001. **b**-**c**, Aggregate profile plots of normalized H2A.Z (b) and H3K9me3 (c) CUT&Tag signals over LTR, tRNA and LINE in 5Aza-dC and TDCIPP treated zebrafish embryos. Repeat lengths scaled into 1kb together with 0.5kb both upstream and downstream were used for aggregate profile plots. **d**, log2FC ofH3K9me3 over quantiles of primed repeats based on log2FC of H2A.Z(5Aza-dC/DMSO). Statistical significance was determined using Student’s t-test, ***, p-value < 0.001. **e**, IGV genome browser snapshots of primed repeats loci for H2A.Z and H3K9me3 following 5Aza-dC exposure.

**Extended Data Fig.6.**
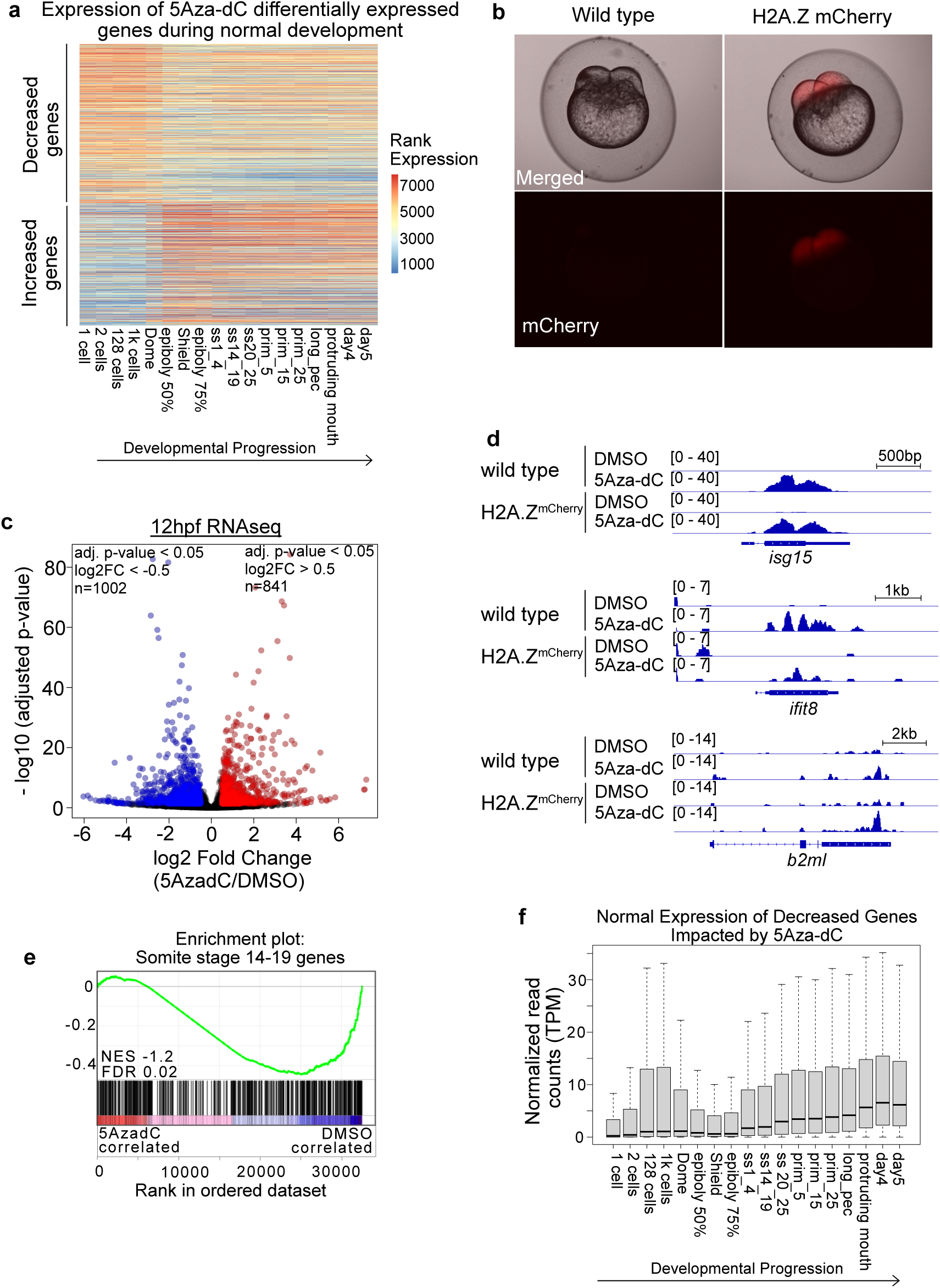
Gene expression signatures of 6hpf and 12hpf wildtype and H2A.Z^mCherry^ embryos after 5Aza-dC exposure. **a**, Heatmap of ranked gene expression levels across developmental stages (from 1-cell stage to day 5) of those decreased genes (n=4,810) and increased genes (n=3,952) in 6hpf embryos after 5Aza-dC exposure. Gene expression read count table (TPM) is from White *et al*, eLife, 2017. **b**, Representative images of wild type and H2A.Z^mCherry^ embryos demonstrating mCherry expression in 2-cell embryos. **c**, Volcano plot showing differential gene expression in 5Aza-dC treated wild type embryos at 12hpf. **d**, Genome browser snapshots of gene expression from 5Aza-dC treated wild type and H2A.Z^mCherry^ embryos at 12hpf. **e**, Gene Set Enrichment Analysis (GSEA) of maximum expressed genes in somite stage 14-19 for transcriptome of 12hpf 5Aza-dC treated wild type embryos. **f**, Boxplot of normalized gene expression read counts (TPM) across developmental stages for 5Aza-dC decreased genes in 12hpf wild type embryos. Gene expression read count table (TPM) is from White *et al*, eLife, 2017.

## Notes

### Competing Interest Statement

The authors have declared no competing interest.

